# Decoding the neural basis of sensory phenotypes in autism

**DOI:** 10.1101/2025.02.27.640592

**Authors:** Matthew Kolisnyk, Kathleen Lyons, Eun Jung Choi, Marlee M. Vandewouw, Bobby Stojanoski, Evdokia Anagnostou, Azadeh Kushki, Rob Nicolson, Elizabeth Kelley, Stelios Georgiades, Jason Lerch, Jennifer Crosbie, Russell Schachar, Muhammad Ayub, Jessica Jones, Paul Arnold, Xudong Liu, Ryan Stevenson

## Abstract

**Background:** Differences in sensory processing are a defining characteristic of autism, affecting up to 87% of autistic individuals. These differences cause widespread perceptual changes that can negatively impact cognition, development, and daily functioning. Recent research identified five sensory processing ‘phenotypes’ with varied behavioural presentations; however, their neural basis remains unclear. This study aims to ground these sensory phenotypes in unique patterns of functional connectivity.

**Methods:** We analyzed data from 146 autistic participants in the Province of Ontario Neurodevelopmental Network. We classified participants into sensory phenotypes using k-means clustering of scores from the Short Sensory Profile. We then computed a connectivity matrix from 200 cortical and 32 subcortical regions and calculated graph-theoretic measures (betweenness centrality, strength, local efficiency, clustering coefficient) to assess information exchange between these regions. We then trained machine learning models to use these measures to classify between all pairs of sensory phenotypes.

**Results:** We replicated that our sample of autistic participants was best categorized into five sensory phenotypes. The machine learning models distinguished 7/10 phenotype pairs using graph-theoretic measures (*p* < 0.005). Information exchange within and between the somatomotor network, orbitofrontal cortex, posterior parietal cortex, prefrontal cortex and subcortical areas were highly predictive of sensory phenotype.

**Conclusions:** This study shows that distinct sensory phenotypes in autism correspond with unique patterns of functional connectivity. Cortical, subcortical, and network-level connectivity all play a role in shaping distinct sensory processing styles in autism. These findings lay the groundwork for understanding these phenotypes and highlight opportunities to develop interventions in cases of maladaptive sensory processing.

## Introduction

Autism is a neurodevelopmental condition characterized by difficulties in communication, social interaction, and restrictive or repetitive behaviours and interests (1) A common feature of autism is differences in sensory processing, with many individuals reporting that they perceive the world differently from their neurotypical peers (2) Sensory processing is fundamental to how we make sense and navigate our world, and as such any differences can profoundly influence perception across a wide range of domains. Indeed, evidence suggests that altered sensory processing can affect high-order cognition (3–5), disrupt developmental trajectories (6,7), and cause significant distress in autistic individuals (8,9).

While sensory differences have been reported in up to 87% of autistic individuals (10), their presentation varies considerably. Individuals may show both hyper- and hypo-sensitivities in sensory processing which may affect all sensory domains or selectively impact one modality (11,12). While this heterogeneity may appear to be unique to each individual, recent research has found reliable patterns of sensory differences across autistic individuals (10,13–15)

A recent large-scale study of autistic children and adolescents (14) identified five distinct sensory processing styles based on scores from the Short Sensory Profile (16). These distinct ‘sensory phenotypes’ were categorized as: 1) sensory adaptive, 2) generalized sensory differences, 3) taste and smell sensitivity, 4) under-responsive and sensation seeking, and 5) movement difficulties with low energy. They represent a spectrum of sensory dysfunction, ranging from minimal (e.g., sensory adaptive), to widespread (e.g., generalized sensory differences), to selective differences within specific domains (e.g., taste and smell sensitivity, under-responsive and sensation seeking, and movement difficulties with low energy). Notably, each phenotype was associated with distinct patterns of social communication, repetitive and restrictive behaviour and interests, and co-occurring symptoms (14,15).

These distinct sensory phenotypes are likely associated with differences in the brain’s functional properties, however, characterizing the distinct neural mechanisms associated with autism remains elusive, and comparisons with neurotypical participants are often conflicting (17,18). For instance, some studies report reduced, while others report enhanced functional connectivity in autism, particularly in the default mode and frontoparietal networks (18–24). One potential explanation is that distinct sensory phenotypes may correspond to unique patterns of brain connectivity, complicating direct comparisons between autistic and neurotypical populations. Investigating this relationship could help clarify the neural mechanisms underlying sensory processing differences and help resolve inconsistencies in connectivity findings within autism research.

To this end, this study sought to investigate how sensory phenotypes in autistic children and adolescents relate to differences in resting-state functional connectivity. Using data from the Province of Ontario Neurodevelopmental Network (POND), which includes resting-state fMRI scans as well as sensory processing scores from autistic participants, we combined graph theory and machine learning to examine functional brain network properties across distinct sensory phenotypes. Overall, to advance our understanding of sensory phenotypes and provide insights into the neural variability underlying these groups. The heterogeneity of autism means that effective supports and treatments for one sensory phenotype may be ineffective in another, underscoring the need to understand the biological mechanisms underlying these phenotypes to effectively establish targeted interventions.

## Methods and Materials

### Data Source

We obtained data for this study from the POND Network (https://pond-network.ca). Research Ethics Boards at each participating institution approved the POND study protocol. All participating institutions obtained written informed consent and assent from the primary caregiver and/or participant.

### Participants

The current study included participants from the POND Network if they had a diagnosis of autism. The study excluded participants if their neuroimaging data contained artifacts (described below) or if their parent or guardian had not completed the Short Sensory Profile. The final dataset contained 146 autistic participants (*Mage* = 10.603, *SD* = 4.400, *Range* = [3,21], 36 female).

### Sensory Phenotype Classification

We used the Short Sensory Profile (16), a 38-item parent-report questionnaire, to measure behaviours associated with different domains of sensory processing. The questionnaire scores participants according to seven different sensory domains: tactile, taste/smell, movement, under-responsive/sensory seeking, auditory filtering, low energy, and visual/auditory. Higher scores reflect less sensory differences, whereas lower scores reflect higher sensory differences. We applied k-means clustering to the Short Sensory Profile scores and identified five different clusters -- corresponding to the five sensory phenotypes previously reported (14,15): (1) sensory adaptive, (2) generalized sensory differences, (3) taste and smell sensitivity, (4) under-responsive and sensation seeking, and (5) movement difficulties with low energy.

### MRI Acquisition

The POND Network collected the MRI data using either a 3T Siemen’s MAGNETOM Trio with a 12-channel head coil or a 3T Siemen’s PrismaFIT with a 20-channel head and neck coil. Participants completed a five-minute T1-weighted image (Trio: TR/TE: 2300/2.96ms; FA: 9°; FOV: 192×240×256mm; 1.0mm isotropic; Prisma: TR/TE: 1870/3.14ms, FA: 9°, FOV: 192×240×256mm, 0.8mm isotropic). This was followed by a five-minute T2* resting state fMRI scan (Trio: TR/TE: 2300/2.96ms; FA: 9°; FOV: 192×240×256mm; 1.0mm isotropic; Prisma: TR/TE: 1870/3.14ms, FA: 9°, FOV: 192×240×256mm, 0.8mm isotropic). During the resting-state fMRI scans, participants viewed either a movie of their choice or Inscapes, a naturalistic movie paradigm designed to be engaging while minimizing cognitive demands (25).

### MRI Preprocessing

We preprocessed the MRI data using fMRIPrep (26), which is built using Nipype (27,28) in Python. For detailed preprocessing steps generated from fMRIPrep, see the supplementary materials.

Briefly, we corrected each participant’s T1-weighted image for intensity non-uniformity, skull-stripped it, and segmented it into gray matter, white matter, and cerebrospinal fluid before being spatially normalizing it to a pediatric template. For the functional data, we applied slice-time correction, motion correction, and susceptibility distortion correction, before we co-registered it to the participant’s T1-weighted image.

Before analysis, we excluded participants with excessive motion artifacts using framewise displacement (FD) and the standardized derivative of root mean square variance over voxels (DVARS). We excluded participants if more than one-third of their frames exceeded a threshold of 0.5 mm FD or 1.5 standardized DVARS. To remove potential confounding signals, we regressed six motion parameters (3 translations and 3 rotations), white matter and cerebrospinal fluid signals, as well as their derivatives and quadratic terms. Finally, we applied high-pass temporal filtering (at 0.008 Hz).

We parcellated the functional data using the Schaefer cortical atlas (29) (200 regions) and the Melbourne subcortical atlas (30) (32 regions), yielding 232 regions of interest. We computed functional connectivity by calculating the Pearson correlation between the time series for each region, resulting in a 232×232 connectivity matrix for each participant. For subsequent graph theory analysis, we binarized these matrices using a threshold of 0.2, which we selected to produce representative networks that are not too sparse to be biologically implausible (31).

### Functional Connectivity Analysis

Graph theory has emerged as an effective analysis method to understand functional connectivity and information exchange between brain regions (32–34). Our analyses focused on four canonical graph theoretic measures to differentiate between sensory phenotypes: betweenness centrality, strength, local efficiency, and clustering coefficient. Briefly, these measures attempt to represent information exchange between different brain regions at a local scale (*i.e.,* local efficiency and clustering coefficient), by a region’s overall connectedness to other regions (*i.e.,* strength), or a region’s ability to act as a junction for information flow between other regions (*i.e.,* betweenness centrality). For the derivation and description of these measures please see the Supplementary Materials. We used MATLAB’s Brain Connectivity Toolbox (35) to estimate each graph measure from each participant’s functional connectivity matrix. This yielded a total of 928 features (232 regions x 4 measures) for subsequent analysis.

### Graph Theory Preprocessing

Before applying machine learning, we preprocessed each graph theory feature. We began by normalizing the distributions of betweenness centrality and strength features using a logarithmic transform. Next, we identified and removed outliers, defined as features exceeding 3 standard deviations, and later interpolated these during the training stages of machine learning. This step resulted in only a small percentage of outliers across all measures (range: 0.345% - 1.104%).

To account for known age-related differences in functional connectivity measures (36–38) and sensory phenotypes (14), we used linear regression to control for age for each feature. We then used this preprocessed dataset, comprising 928 features across 146 participants, to build models that distinguish between each pair of sensory phenotypes.

### Predicting Sensory Phenotypes

We conducted the machine learning analyses on Python (version 3.11.5) using the sci-kit learn package for model fitting and the hyperopt package for hyperparameter tuning (39,40). If the graph theory features contain useful information to distinguish between different sensory phenotypes, then machine learning should be able to distinguish between all pairs of sensory phenotypes. We stratified the data for each comparison into three pseudorandom folds, with the requirement that an equal ratio of each sensory phenotype was represented in both the training (model fitting) and test (model prediction) sets. Within the training data, we performed an additional three-fold stratification for hyperparameter optimization.

We implemented the following training approach: first, we imputed outliers in the training set using K-Nearest Neighbor imputation. Next, we standardized each graph-theory measure to have a mean of 0 and a standard deviation of 1. We then fit a linear Support Vector Classifier (SVC) to the data (41). Due to the large number of features, we regularized the SVC using the hyperparameters: *C*, number of iterations and tolerance, which we optimized using a Bayesian optimization framework (39). The objective of the optimization was to maximize balanced accuracy (on the training set), defined as the arithmetic mean of sensitivity and specificity. Importantly, this optimization process had no access to the data in the test set.

We applied the optimized model to the test dataset and evaluated the classification performance using balanced accuracy. To measure the importance of each feature, we extracted the standardized coefficients of the linear SVC. We repeated the training and testing procedure ten times to assess performance across different stratified samples of the data. Finally, we averaged the balanced accuracy scores and coefficients across the three stratified folds and repeated these procedures ten times (each with a different train-test split) to obtain robust estimates of model performance and feature importance.

## Statistical Analysis

### Sensory Phenotypes Performance Metrics

We used permutation testing to assess the statistical significance of the classification performance. This approach is well-suited for machine learning analyses as it accurately measures true chance-level performance and accounts for potential idiosyncrasies in the data that could drive above-chance results (42,43).

For each of the ten sensory phenotype comparisons, we generated a null hypothesis by simulating chance-level balanced accuracy. Specifically, we applied the same machine learning approach described above but randomized the sensory phenotype labels before training the model. We repeated this procedure for 1000 permutations to create a distribution of balanced accuracy scores. We then calculated *p*-values by comparing the true balanced accuracy to the null distribution. To account for multiple comparisons, we applied Bonferroni correction, considering results significant at *p* < 0.005.

### Determining Influential Graph Theory Predictors of Sensory Phenotypes

We applied a similar permutation testing approach to identify significant graph theory measures and regions that contributed to accurate phenotype decoding. For each comparison, we extracted standardized coefficients from the linear SVC model across folds and iterations. We repeated this process for 1000 permutations to determine the range of coefficients that could arise by chance.

To correct for multiple comparisons, we used the max-*t* method (43). Specifically, we calculated the maximum and minimum standardized coefficients across features and iterations for each permutation. Over 1000 permutations, we generated two distributions representing the maximum and minimum coefficients obtainable by chance. We identified highly influential predictors as those whose coefficients exceeded the 97.5th percentile or fell below the 2.5th percentile for the maximum and minimum distribution, respectively.

### Graph Theory Measures and Network-Level Analysis

In addition to evaluating individual features, we examined how coefficients varied within graph theory measures (e.g., clustering coefficient) and brain networks. Networks included the Default Mode, Dorsal Attention, Frontoparietal, Ventral Attention, Limbic, Somatomotor, and Visual networks, as well as the subcortical parcellation, which we treated as an additional ‘network’.

For each sensory phenotype comparison, we calculated the absolute value of coefficients split by phenotype and averaged them within each graph theory measure or network. We then conducted Welch’s *t-*tests to compare coefficients across sensory phenotype pairs. To correct for multiple comparisons, we used the Benjamini-Hochberg method.

## Results

### Decomposing Sensory Scores into Sensory Phenotypes

Replicating previous results (14,15), the Short Sensory Profile scores were best decomposed into five clusters of distinct sensory phenotypes (see Figure 1A). The first phenotype, sensory adaptive (SA), is characterized by high (smaller sensory differences) scores across all sensory domains. The second, generalized sensory differences (GSD), displays low (larger sensory differences) scores across all domains. The third phenotype, taste and smell sensitivity (TSS), is defined by selectively low scores in the taste and smell domain. The fourth phenotype, under-responsive and sensation seeking (URSS), exhibits relatively high scores in most domains, with the exception of the under-responsive domain. Lastly, movement difficulties with low energy (MDLE) is marked by low scores in the movement and low energy domains. Demographic information for each group is presented in Table 1.

**Figure 1.**
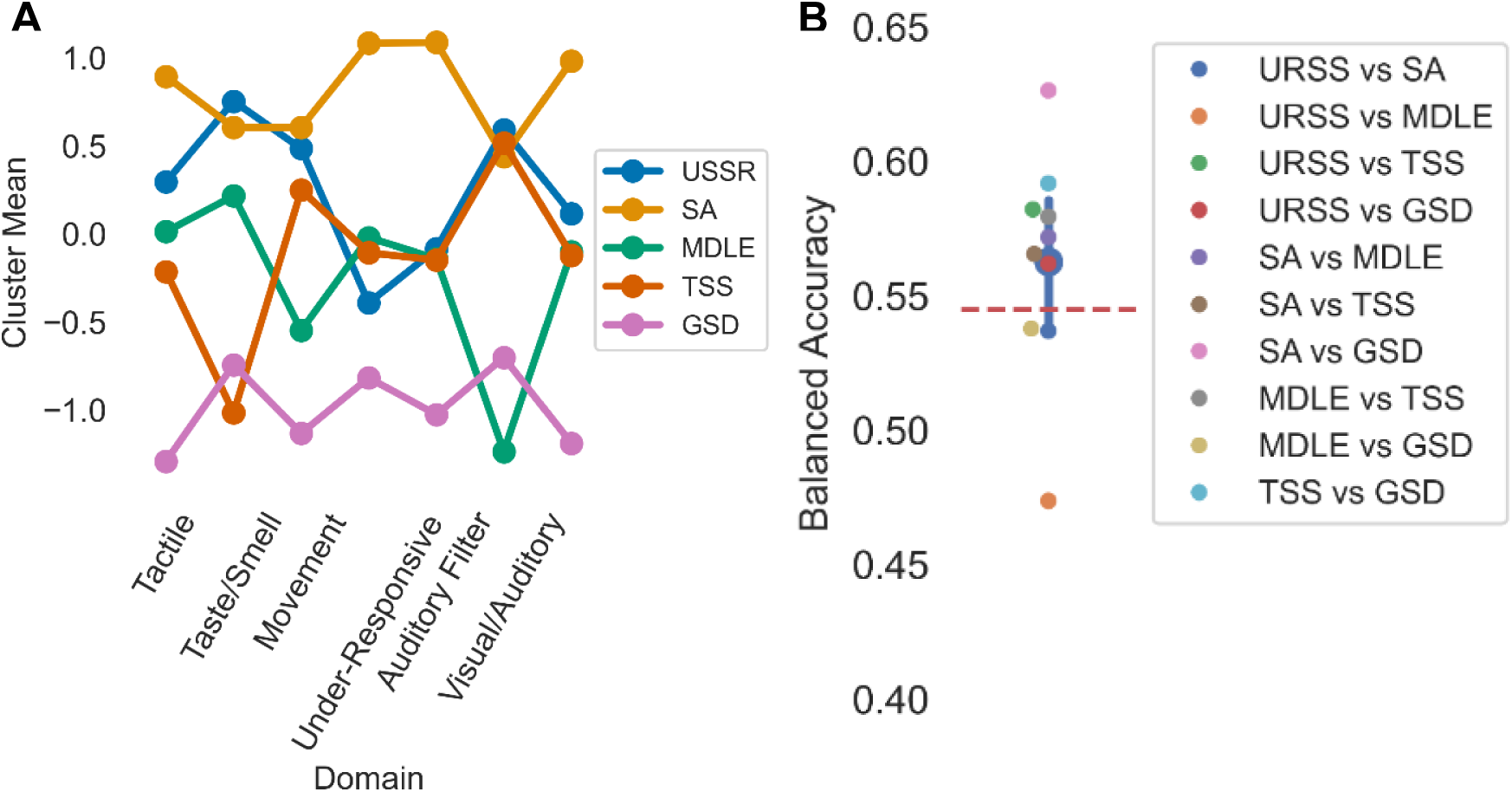
(A) Line plot illustrating how the different sensory phenotypes align with the domains of the short sensory profile. (B) Point plot showing the balanced accuracy score computed for each sensory phenotype pair. The red dashed line separates significant comparisons (p < 0.005) from non-significant ones. The large blue point represents the overall average balanced accuracy score across all phenotype comparisons, with error bars indicating the 95% confidence interval. This overall score was significantly above chance (see Supplementary Materials). Abbreviations: sensory adaptive (SA), generalized sensory differences (GSD), taste and smell sensitivity (TSS), under-responsive and sensation seeking (URSS), movement difficulties with low energy (MDLE).

**Table 1.** Demographic and Sample Information for Sensory Phenotypes.

| **Sensory Phenotype** | ***n*** | **Age (Mean, SD)** | **Sex** |
| --- | --- | --- | --- |
| Sensory adaptive | 39 | 11.205 (5.100) | 27 Male; 12 Female |
| Generalized sensory differences | 26 | 9.692 (4.145) | 22 Male; 4 Female |
| Taste and smell sensitivity | 22 | 10.455 (4.522) | 13 Male; 9 Female |
| Under-responsive and sensation seeking | 29 | 9.621 (4.059) | 24 Male; 5 Female |
| Movement difficulties with low energy | 30 | 11.667 (3.698) | 24 Male; 6 Female |

### Decoding Sensory Phenotypes

Seven of the ten sensory phenotype pairs were significantly distinguished by the graph theoretic measures (see Figure 1B). This included: URSS vs TSS (Balanced Accuracy = .582, *p* < .001), URSS vs GSD (Balanced Accuracy = .562, *p =* 0.002), SA vs MDLE (Balanced Accuracy = .572, *p* < .001), SA vs TSS (Balanced Accuracy = .566, *p =* .002), SA vs GSD (Balanced Accuracy = .626, *p* < .001), MDLE vs TSS (Balanced Accuracy = .579, *p* < .001), and TSS vs GSD (Balanced Accuracy = .592, *p* < .001). In contrast, three comparisons -- URSS vs. GSD, URSS vs. SA, and MDLE vs. GSD -- did not yield balanced accuracy scores significantly greater than chance (all p > 0.025).

To better understand the decoding results, we conducted two post-hoc analyses (see the Supplementary Materials). Briefly, the first analysis revealed that the average balanced accuracy score across all pairwise comparisons was significantly greater than chance, providing support that these sensory phenotypes produce distinct functional connectivity profiles. The second analysis suggested that decoding performance was stronger for phenotype pairs with greater dissimilarity in their cluster mean scores, provided some evidence that the distinctiveness of graph measures seems to be influenced by intrinsic sensory differences between phenotypes.

## Neural Basis of Decoded Sensory Phenotypes

### Sensory Adaptive vs. Generalized Sensory Differences

The comparison between SA and GSD yielded the highest decoding performance of all pairs, likely because the underlying clusters reflect the extremes of sensory processing (i.e., minimal vs. widespread differences). As shown in Figure 2A, several regions were significant predictors of the differences between these groups.

**Figure 2.**
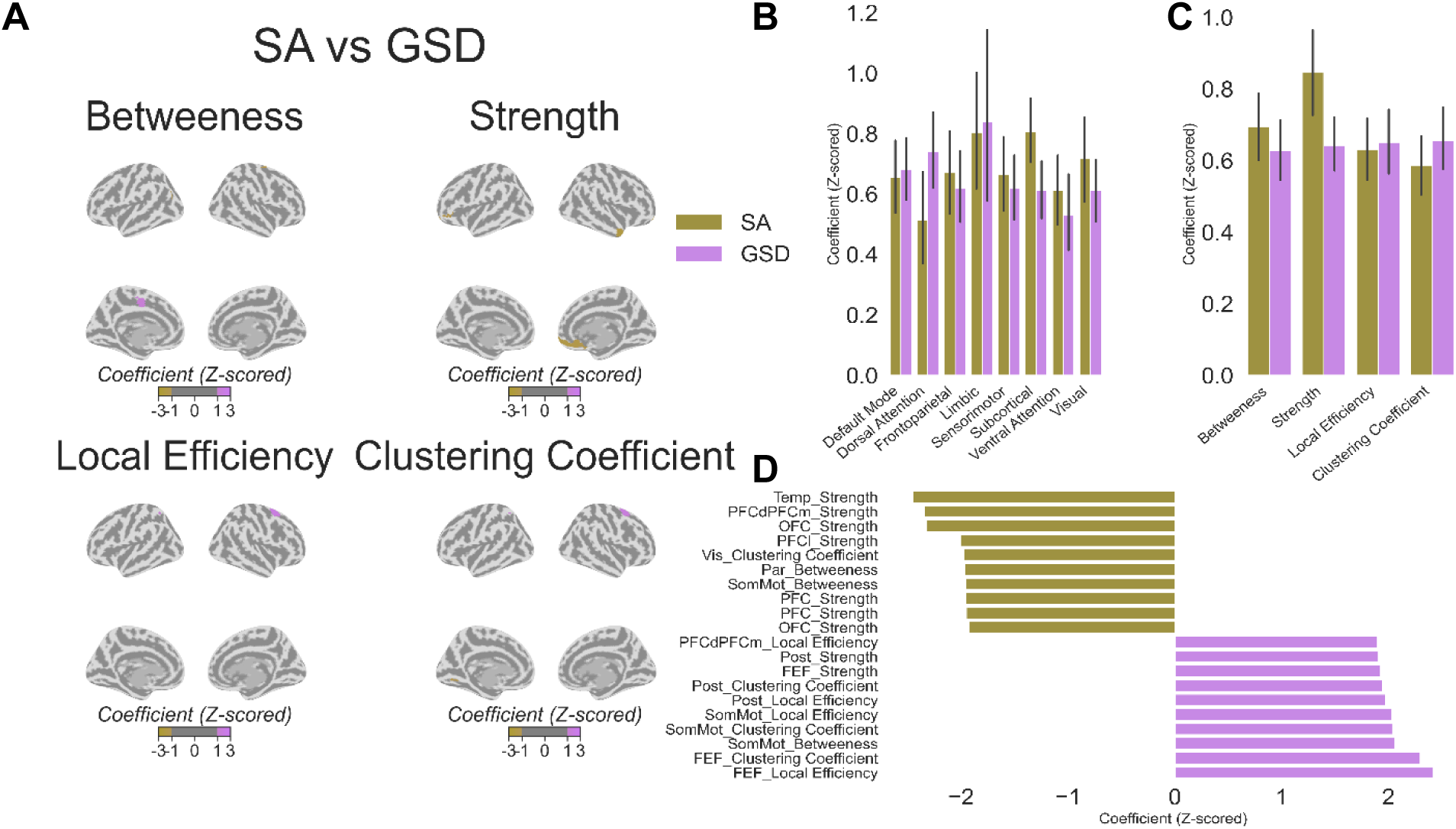
(A) Surface map illustrating statistically significant coefficients (converted to *Z*-scores) for the sensory adaptive (gold) and generalized sensory differences (magenta) phenotypes. (B) Bar plots displaying the coefficients averaged across each network for both phenotypes, with vertical lines representing 95% confidence intervals. (C) Bar plots showing the coefficient averaged across each graph theory measure for both phenotypes, with vertical lines representing 95% confidence intervals. (D) Bar plots highlighting the top ten regions by graph theory measure combination with the largest coefficient for each phenotype. Abbreviations: sensory adaptive (SA); generalized sensory differences (GSD); Temporal (Temp); Dorsal Prefrontal Cortex (PFCd); Medial Prefrontal Cortex (PFCm); Orbitofrontal Cortex (OFC); Lateral Prefrontal Cortex (PFCl); Posterior (Post); Frontal Eye Fields (FEF); Somatomotor (SomMot).

Broadly, the SA group was characterized by larger betweenness and strength features, particularly within the default mode network. In contrast, local efficiency and clustering coefficient, particularly within the dorsal attention network, were more predictive of the GSD group. In addition, the SA group showed larger betweenness in the posterior parietal and somatosensory cortex, as well as greater strength in the temporal lobe, orbitofrontal cortex, and lateral and medial prefrontal cortex. Conversely, the GSD group demonstrated larger local efficiency and clustering coefficients in the posterior parietal and frontal eye fields. Supplementary Table 1 provides a complete list of significant predictors.

When we collapsed the coefficients across networks (Figure 2B) and graph theory measures (Figure 2C), no comparisons survived correction. However, the subcortical network showed a trend toward significance (*t*(124.692) = 2.684, *p =* 0.008), with coefficients being, on average, larger in the SA group.

### Sensory Adaptive vs Movement Difficulties with Low Energy

Several regions and graph measures allowed for effective decoding between SA and MDLE groups (see Figure 3A). Many regions in the parietal cortex (e.g., posterior parietal cortex, postcentral gyrus) had larger scores in the MDLE group for most graph-theory measures, except for one posterior parietal region where the SA group showed higher betweenness. Betweenness scores were also higher in the parahippocampal gyrus and temporal pole for the MDLE group.

**Figure 3.**
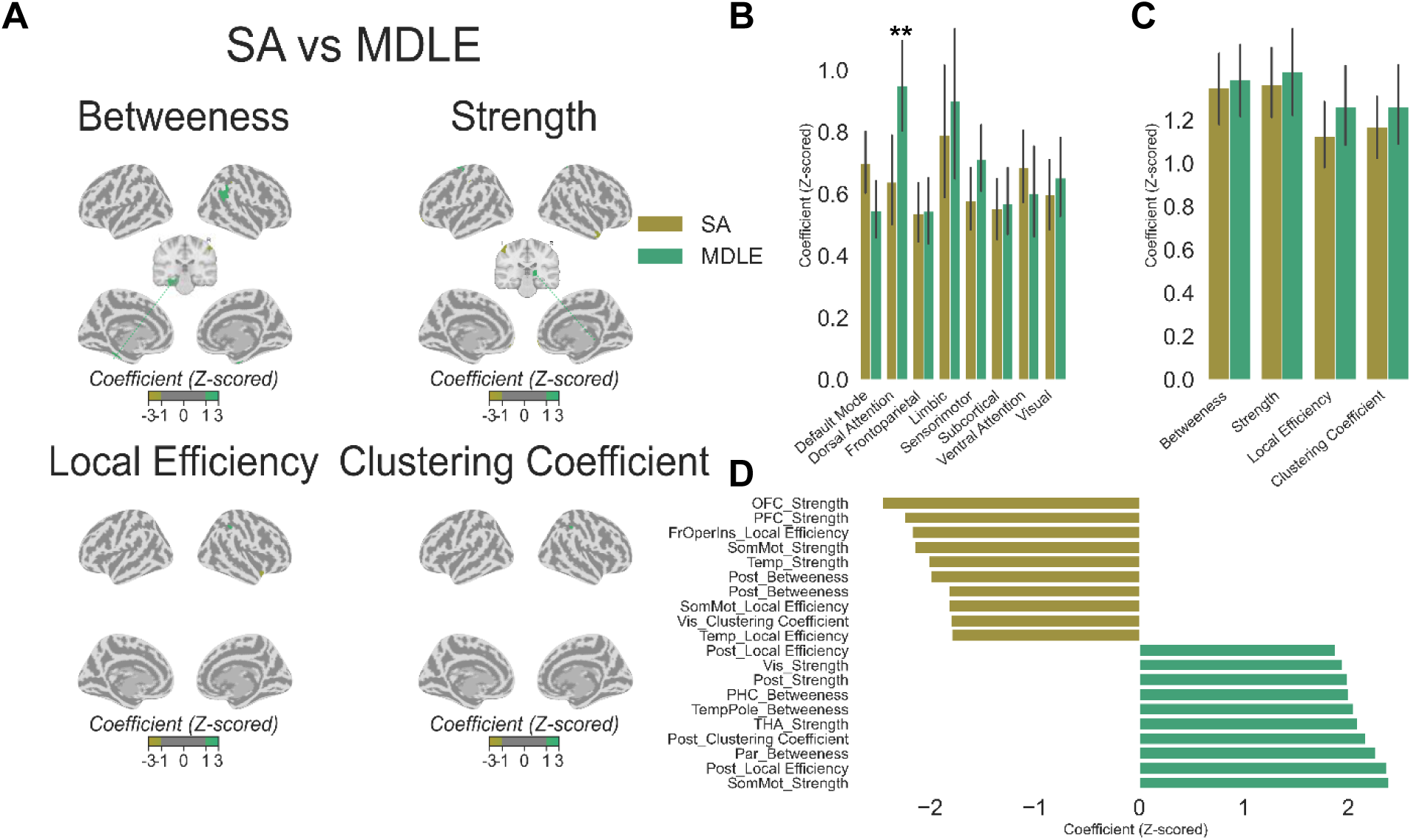
(A) Surface map illustrating statistically significant coefficients (converted to *Z*-scores) for the sensory adaptive (gold) and movement difficulties with low energy (green) phenotypes. Inset in the centre of the surface maps contain a horizontal slice to highlight relevant medial regions (the parahippocampal gyrus (for betweenness) and thalamus (for strength) coordinates = 0, -28, 12). (B) Bar plots displaying the coefficients averaged across each network for both phenotypes, with vertical lines representing 95% confidence intervals. (C) Bar plots showing the coefficient averaged across each graph theory measure for both phenotypes, with vertical lines representing 95% confidence intervals. (D) Bar plots highlighting the top ten regions by graph theory measure combination with the largest coefficient for each phenotype. Abbreviations: sensory adaptive (SA); movement difficulties and low energy (MDLE). Temporal (Temp); Orbitofrontal Cortex (OFC); Prefrontal Cortex (PFC); Posterior (Post); Parietal (Par); Frontal Eye Fields (FEF); Somatomotor (SomMot); Visual Cortex (Vis); Frontal Operculum Insula (FrOperIns); Parahippocampal Cortex (PHC); Thalamus (THA).

Strength scores differed between groups in several regions. The SA group had higher strength in the orbitofrontal cortex, prefrontal cortex, and temporal lobe, while the MDLE group showed higher strength in the dorsoposterior thalamus. In the somatosensory cortex, strength varied by region, with some regions linked to SA and others to MDLE. Supplementary Table 2 provides a complete list of significant features.

When we averaged coefficients across networks (Figure 3B), the dorsal attention network had significantly larger coefficients in the MDLE group (*t*(93.616) = -2.931, *p =* 0.004). When coefficients were averaged across graph-theory measures (Figure 3C), no comparisons survived correction.

### Under-responsive and sensation seeking vs generalized sensory differences

The GSD group generally showed regions with larger local efficiency and clustering coefficient, while the URSS group showed specific regions with larger betweenness and strength. For the GSD group, local efficiency was higher in the medial prefrontal cortex, and clustering coefficients were larger in the somatosensory cortex, as well as in two subcortical regions: the ventroposterior thalamus and nucleus accumbens core.

In contrast, the URSS group demonstrated larger betweenness in the visual cortex, medial frontal cortex, lateral prefrontal cortex, and precuneus, along with greater strength in the orbitofrontal cortex. See Figure 4A for a surface map of these results and Supplementary Table 3 for a complete list of significant predictors. When coefficients were averaged across networks (Figure 4B) and graph-theory measures (Figure 4C), no comparisons survived correction.

**Figure 4.**
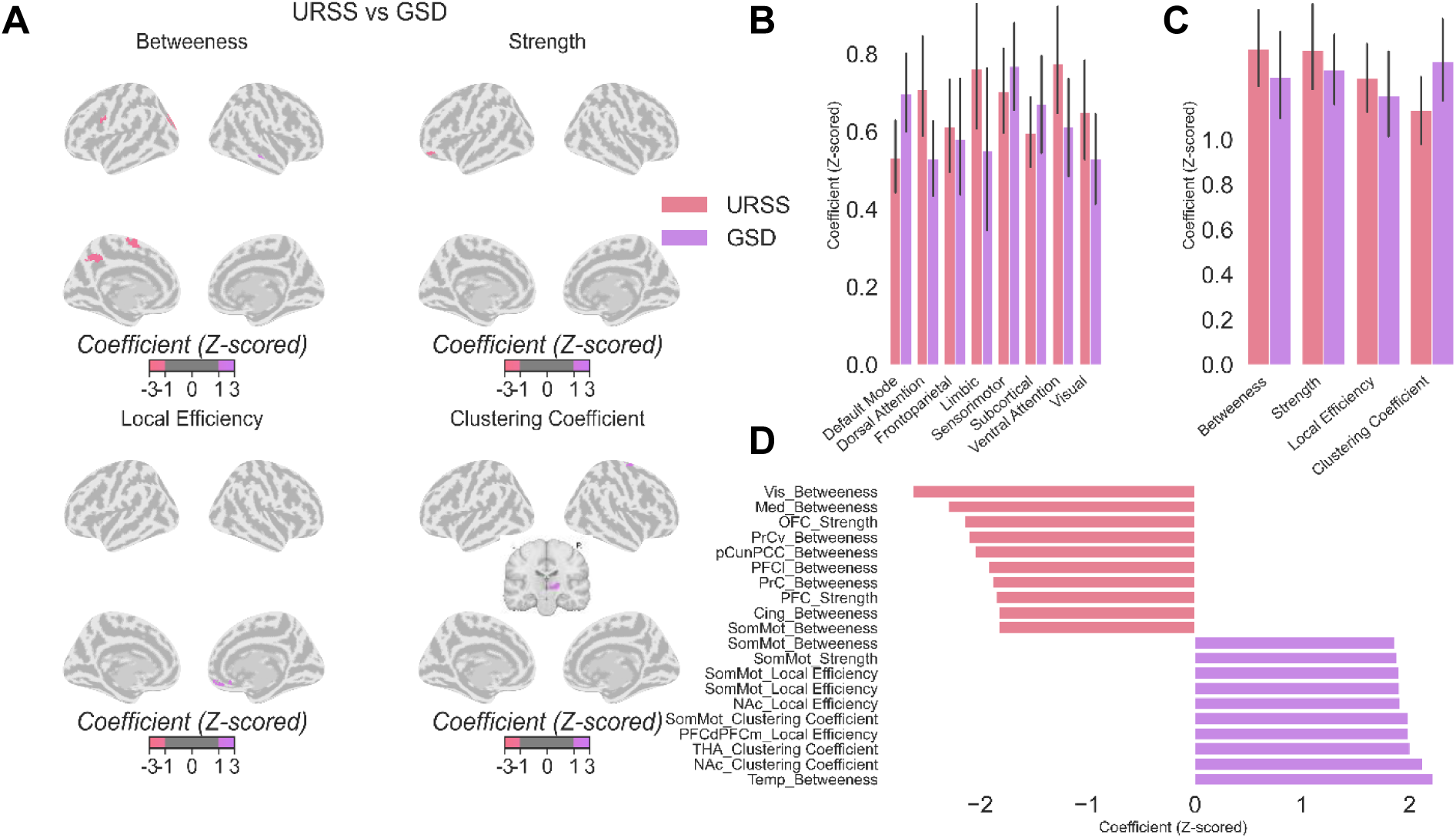
(A) Surface map illustrating statistically significant coefficients (converted to *Z*-scores) for the under-responsive and sensation seeking (red) and generalized sensory difficulties (magenta) phenotypes. Inset in the centre of the surface maps contain a horizontal slice to highlight relevant medial regions (the thalamus and nucleus accumbens (for clustering coefficient) coordinates = 0, -20, -12). (B) Bar plots displaying the coefficients averaged across each network for both phenotypes, with vertical lines representing 95% confidence intervals. (C) Bar plots showing the coefficient averaged across each graph theory measure for both phenotypes, with vertical lines representing 95% confidence intervals. (D) Bar plots highlighting the top ten regions by graph theory measure combination with the largest coefficient for each phenotype. Abbreviations: under-responsive and sensation seeking (URSS); generalized sensory difficulties (GSD); Cingulum (Cing); Nucleus Accumbens Core (NAc); Medial (Med); Precentral Ventral (PrCv); Precentral (PrC); Temporal (Temp); Orbitofrontal Cortex (OFC); Precuneus Posterior Cingulate (pCunPCC); Medial Prefrontal Cortex (PFCm); Lateral Prefrontal Cortex (PFCl); Prefrontal Cortex (PFC); Posterior (Post); Somatomotor (SomMot); Thalamus (THA).

### Connectivity Profiles of Taste and Smell Sensitivity

The TSS phenotype showed distinct connectivity profiles that differentiated it from all other sensory phenotypes. Across comparisons, TSS consistently exhibited larger betweenness in the posterior parietal cortex. Additionally, either strength or betweenness in the orbitofrontal cortex and somatosensory cortex were key features distinguishing TSS from other phenotypes. Betweenness was almost always larger in the precuneus for TSS, whereas other phenotypes showed larger betweenness in the medial prefrontal cortex and visual cortex. Local efficiency and clustering coefficient in the medial prefrontal cortex were generally higher for TSS compared to other phenotypes. However, regions within the somatosensory cortex typically showed lower clustering coefficient and local efficiency for TSS, except when compared to the GSD phenotype.

When we averaged the coefficients across networks, TSS showed larger coefficients in the dorsal attention (*t*(76.760) = 2.817, *p =* 0.006) and ventral attention networks (*t*(85.959) = 2.986, *p =* 0.004), but smaller coefficients in subcortical regions (*t*(84.392) = -3.136, *p =* 0.002) compared to the SA group. TSS had smaller coefficients in the frontoparietal network (*t*(116.649) = 2.872, *p =* 0.005) than the URSS group.

Lastly, TSS exhibited smaller coefficients in the subcortical (*t*(114.862) = 3.238, *p =* 0.002) and visual networks (*t*(112.159) = 2.959, *p =* 0.004) compared to the MDLE group. No significant differences were observed when coefficients were averaged across graph-theory measures for any group comparison. See Figure 5 for the comparison between SA and TSS, and the Supplementary Materials for comparisons with other phenotypes. Complete lists of significant features are provided in Supplementary Tables 4S–7S.

**Figure 5.**
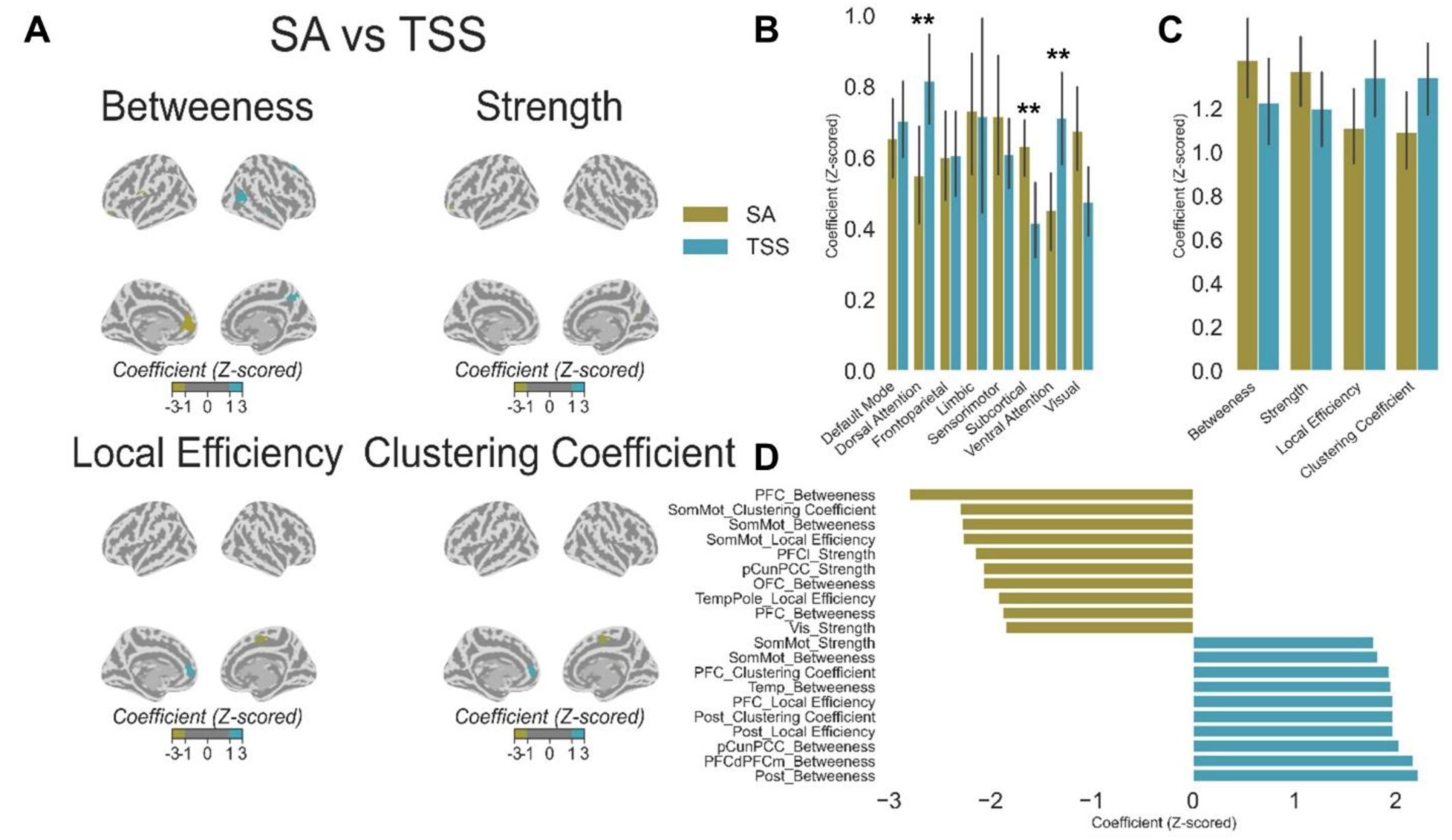
(A) Surface map illustrating statistically significant coefficients (converted to *Z*-scores) for the sensory adaptive (gold) and taste and smell sensitivity (blue) phenotypes. (B) Bar plots displaying the coefficients averaged across each network for both phenotypes, with vertical lines representing 95% confidence intervals. (C) Bar plots showing the coefficient averaged across each graph theory measure for both phenotypes, with vertical lines representing 95% confidence intervals. (D) Bar plots highlighting the top ten regions by graph theory measure combination with the largest coefficient for each phenotype. Abbreviations: sensory adaptive (SA); taste and smell sensitivity (TSS); Temporal (Temp); Orbitofrontal Cortex (OFC); Precuneus Posterior Cingulate (pCunPCC); Medial Prefrontal Cortex (PFCm); Lateral Prefrontal Cortex (PFCl); Prefrontal Cortex (PFC); Posterior (Post); Somatomotor (SomMot).

## Discussion

This study investigated whether distinct sensory phenotypes in autistic children and adolescents could be differentiated based on their resting-state functional connectivity. By integrating machine learning with graph-theory, we aimed to determine whether functional connectivity patterns align with specific sensory profiles previously identified in autism (14,15). Our findings demonstrate that sensory phenotypes in autism can be reliably distinguished based on functional connectivity. Decoding accuracy appeared to be partially driven by intrinsic differences in sensory processing, with the most distinct phenotypes yielding the largest decoding accuracies. The ability to decode these phenotypes reinforces the notion that sensory profiles in autism are not only behaviourally distinct but also reflect meaningful differences in neural connectivity.

Several regional connectivity patterns emerged as consistent predictors across phenotypes, particularly within the somatomotor network, parietal cortex, orbitofrontal cortex, and prefrontal cortex. These regions span the sensory processing hierarchy, encompassing primary and secondary sensory areas, association cortices, and regions involved in decision-making and executive function. Notably, many of these regions have been implicated in studies comparing autistic individuals with neurotypical controls (44–53). Our results further existing research by demonstrating that connectivity profiles in these regions not only differ between autistic and neurotypical individuals but also vary across distinct sensory processing phenotypes within autism.

Differences in attentional networks, such as the dorsal attention and ventral attention, in addition to regions within the default mode network (e.g., the medial prefrontal cortex) helped distinguish between several different phenotypes. Both attentional networks and the default mode network are well known to modulate sensory input and attention and are widely implicated in autism research (18–23). Indeed, a previous study investigating functional connectivity in a large sample of autistic patients (*n* = 796 across 3 datasets) found patterns of both reduced connectivity (within the somatosensory network and between this network and the salience, dorsal attention and visual networks) and increased connectivity (in the default mode network compared to the rest of the brain and between cortical and subcortical networks) (46). While not linked to distinct sensory phenotypes per se, large scale differences in networks have the potential to influence sensory processing by prioritizing or allocating attention resources to specific types of sensory stimuli. Indeed, the involvement of the networks found in the study suggest both top-down control and attentional mechanisms could be critical for the development of distinct sensory processing styles in autism.

Subcortical regions also played a significant role in distinguishing between sensory phenotypes. Specifically, GSD and TSS phenotypes exhibited lower subcortical coefficients compared to the SA group. Additional comparisons, including MDLE vs. TSS, SA vs. MDLE, and URSS vs. GSD, identified subcortical regions as significant predictors of sensory phenotypes. These findings may reflect the canonical role subcortical regions play in sensory gating, processing, and integrating sensory information before it reaches the cortex. Previous studies have noted differences in specific subcortical areas (*i.e.,* inferior colliculus) between typically developing and autistic individuals in speech tasks (54) as well as a disproportionate influence of subcortical regions on sensory regions (55). In addition, these focal subcortical differences are accompanied by network-level alterations between autistic and typically-developing cohorts (46,56). Together, these findings suggest that specific sensory phenotypes may be disproportionately influenced by subcortical dynamics. Future research should investigate how causal relationships between subcortical, regional, and network-level processes constitute distinct sensory processing.

## Limitations

First, not all sensory phenotypes could be decoded with statistical significance, indicating that some phenotypes may overlap or that the sample size may not be sufficient to detect subtle differences in connectivity. Second, the lack of a generalization set limits the ability to validate the findings across broader autistic populations. Third, motion-related confounds have recently been implicated in explaining neural differences between autistic and typically-developing populations (57). However, our findings look at differences across autistic individuals and hence do not have the same concerns. While some phenotypes may be disproportionately affected by these confounds, we removed highly contaminated samples, and regressed motion confounds from the preprocessed fMRI data. Finally, although the Short Sensory Profile is a well-validated tool to categorize sensory phenotypes, its reliance on caregiver-report data introduces subjectivity. Incorporating objective behavioural or neural assessments of sensory responses may improve the accuracy and generalizability of later classifications.

## Conclusion

This study supports the existence of distinct sensory phenotypes in autistic populations and reveals their neural correlates through functional connectivity patterns. These findings suggest that while sensory processing differences in autism are behaviourally diverse, they correspond to systematic variations in neural connectivity. Understanding these neural signatures provides a foundation for recognizing sensory processing in autism as a complex neural phenomenon. Research can continue to clarify the heterogeneity within autism, which may help guide the development of more effective interventions for maladaptive sensory processing.

## Acknowledgments

The Province of Ontario Neurodevelopmental Network is funded through the support of the Ontario Brain Institute, which is funded partially by the Ontario Provincial Government (POND, PIs: EA and JL). This work was also funded through a CIHR Project Grant (487850, PIs: RAS, BS, EA, and AK). RAS is funded through a Dorothy Killam Fellowship, an NSERC Discovery Grant (RGPIN-2024-06233), two SSHRC Insight Grants (435-2017-0936 & 435-2024-1375), a CIHR Project Grant (487850), the University of Western Ontario Faculty Development Research Fund, and a Canadian Foundation for Innovation John R. Evans Leaders Fund (37497), and through a grant from the Canada First Research Excellence Fund (BrainsCAN).

