## Supplementals for "Decoding the neural basis of sensory phenotypes in autism"

### Supplementary Materials

### Neuroimaging preprocessing using fMRIPrep

Neuroimaging data was preprocessed using *fMRIPrep* (Esteban, Markiewicz, et al., 2019; Esteban, Markiewicz, et al., 2019); which is based on *Nipype* (Gorgolewski et al., 2011; Gorgolewski et al., 2018).

Each participant's T1-weighted image was corrected for intensity non-uniformity (INU) with N4BiasFieldCorrection (Tustison et al. 2010) and was used as T1w-reference throughout the workflow. The T1w-reference was then skull-stripped using the OASIS template with Advance Normalization Tools (ANTs; Avants et al. 2008) to generate a brain mask. FreeSurfer (Dale, Fischl, and Sereno 1999) was used to reconstruct brain surfaces, and the brain mask estimated previously was refined with a custom variation of the method to reconcile ANTs-derived and FreeSurfer-derived segmentations of the cortical gray-matter of Mindboggle (Klein et al. 2017). ANTs nonlinear registration was used spatially normalize the brain-extracted image to a pediatric template (Fonov et al., 2009, 2011). FMRIB's Software Library (FSL) was used to segment the brain-extracted normalized image into cerebrospinal fluid (CSF), white matter (WM), and gray matter (Zhang, Brady, and Smith 2001).

Each participant's functional data was slice-time and motion corrected using Analysis of Functional NeuroImages (AFNI23) and FSL24 software, respectively. Fieldmap-less distortion correction was performed by co-registering the data to the corresponding T1-weighted image with intensity inverted (Huntenburg et al., 2014; Wang et al., 2017), constrained with an average fieldmap template (Treiber et al., 2016), implemented with ANTs. The resulting data were co-registered to the T1-weighted image with boundary-based registration using bbregister (FreeSurfer) with six degrees of freedom (Greve and Fischl, 2009). The motion-correcting, functional-to-anatomical, and anatomical-to-template transformations were concatenated and applied in a single step with ANTs using Lanczos interpolation (Lanczos, 1964). Framewise displacement (FD) and the standardized derivative of root mean square variance over voxels (DVARS) was computed using the implementation in Nipype (Power et al. 2014). Participants with more than 1/3 of their frames exceeding a threshold of 0.5 mm FD or 1.5 standardised DVARS were excluded.

The six motion parameters, WM and CSF signals and their derivatives and quadratic terms (Satterthwaite et al. 2013) were regressed from the functional data while performing high-pass temporal filtering (0.008 Hz) using AFNI.

### Calculating Graph Theory Measures

#### *Strength*

In an unweighted network, the strength of a region is simply the number of directly connected regions.

$$S(v) = n_v$$

Where $n_v$ is the number of regions connected to region *v*.

#### *Local Efficiency*

Local efficiency measures how efficiently information is exchanged among locally connected regions after removing a given region (*v)*.

$$LE(v) = \frac{1}{(n_v(n_v - 1))} \sum_{i \neq j} \frac{1}{d_{i,j}}$$

Where $n_v$ is the number of regions locally connected to region *v* and $d_{i,j}$ represents the shortest path length between all pairs of regions (*i* and *j*) which are locally connected to region *v*.

#### *Clustering Coefficient*

The clustering coefficient measures the likelihood that a region's locally connected regions are also connected to each other.

$$C(v) = \frac{2E_v}{(n_v(n_v - 1))}$$

Where $n_v$ is the total number of local connections with region *v* and $E_v$ is the total number of connections between the local connections.

#### *Betweenness Centrality*

Betweenness centrality quantifies the importance of a region as a junction between other connected regions. It is measured by the ratio of how often one region lies on the shortest path compared to all other shortest paths of the connected regions.

$$BC(v) = \sum \frac{\sigma(s, t|v)}{\sigma(s, t)}$$

Where $\sigma(s, t|v)$ is the number of shortest paths from region *s* to region *t* that pass-through region *v* and $\sigma(s, t)$ is the total number of shortest paths from region *s* to region *t*.

### Assessing the Validity of Sensory Phenotypes

We conducted two follow up analyses to further validate the observed decoding performance. The first analysis explored whether the *mean* balanced accuracy score across comparison was also greater than what is expected by chance. If so, this would help validate that, across comparisons, graph theory measures can be used to decode between sensory phenotypes.

Second, to help interpret why the decoding performance of some comparisons were higher than others, we conducted a follow up analysis by correlating the balanced accuracy results for each comparison and an underlying ‘dissimilarity’ between the clusters that define each phenotype. The idea being that more inherently dissimilar clusters should be easier for the machine learning approach to decode. To construct a dissimilarity score, we took the Euclidean distance between the loadings on each short sensory profile test for each phenotype.

The first exploratory analysis revealed that the average balanced score was well above chance (Balanced Accuracy = .563, $p < .001$), further verifying the validity of these distinct sensory phenotypes (see Supplementary Figure 1A). Second, despite there only being 10 observations, we observed a marginally significant correlation between the balanced accuracy between a given comparison and the distance between the loadings of the sensory phenotypes that define that comparison ($r(9) = .570$, $p = .085$; see Supplementary Figure 1B).

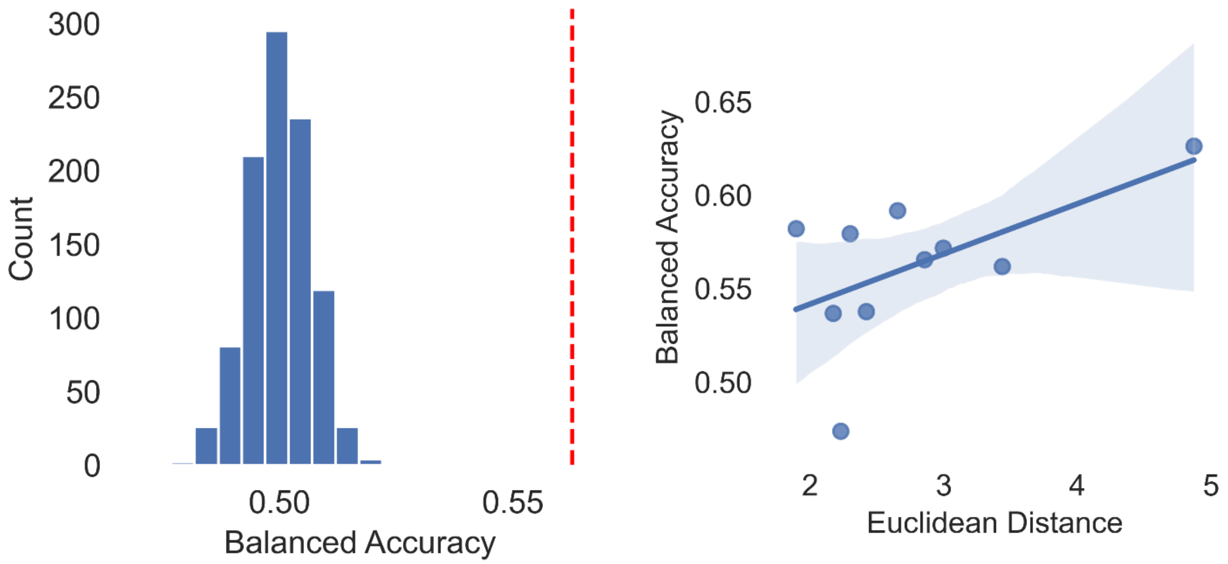


**Supplementary Figure 1**. (A) Histogram of the average balanced accuracy scores expected by chance (via permutation testing). The red dashed line is the actual averaged balanced accuracy score. (B) Scatter plot of the Euclidean distance between phenotype clusters and balanced accuracy scores for a given comparison, including the line of best fit and confidence interval (shaded blue).

### Taste and scent sensitivity additional comparisons

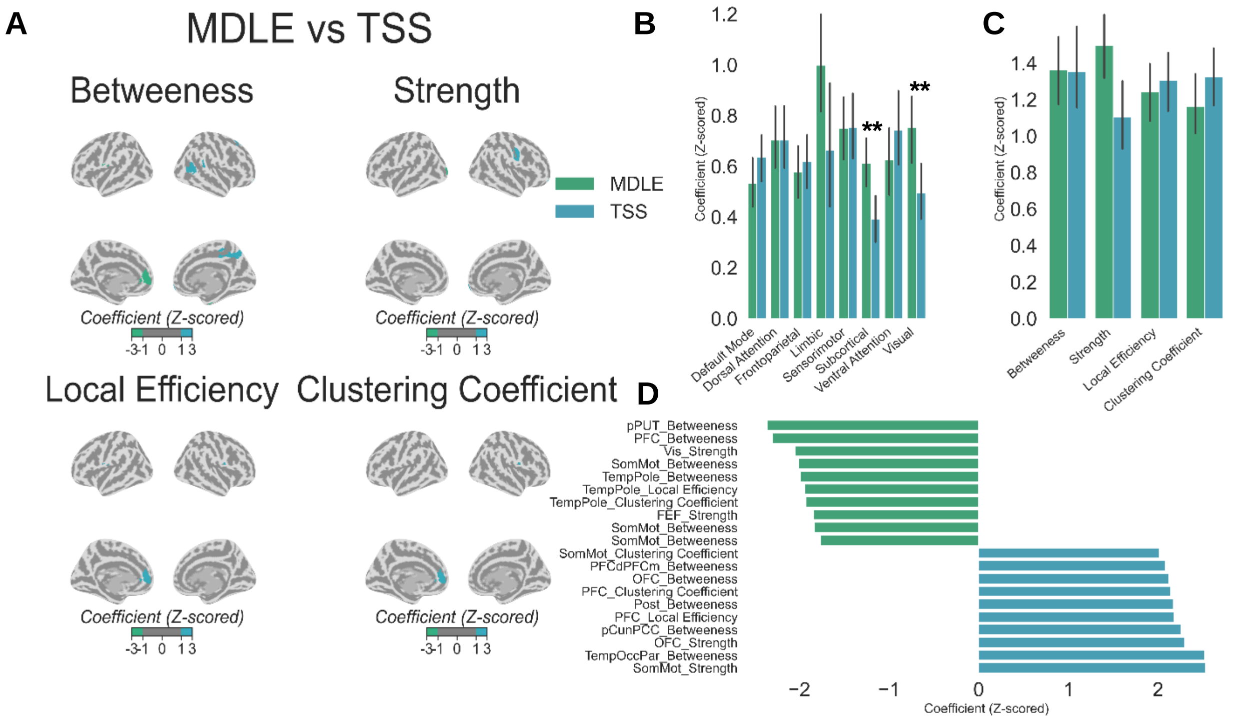


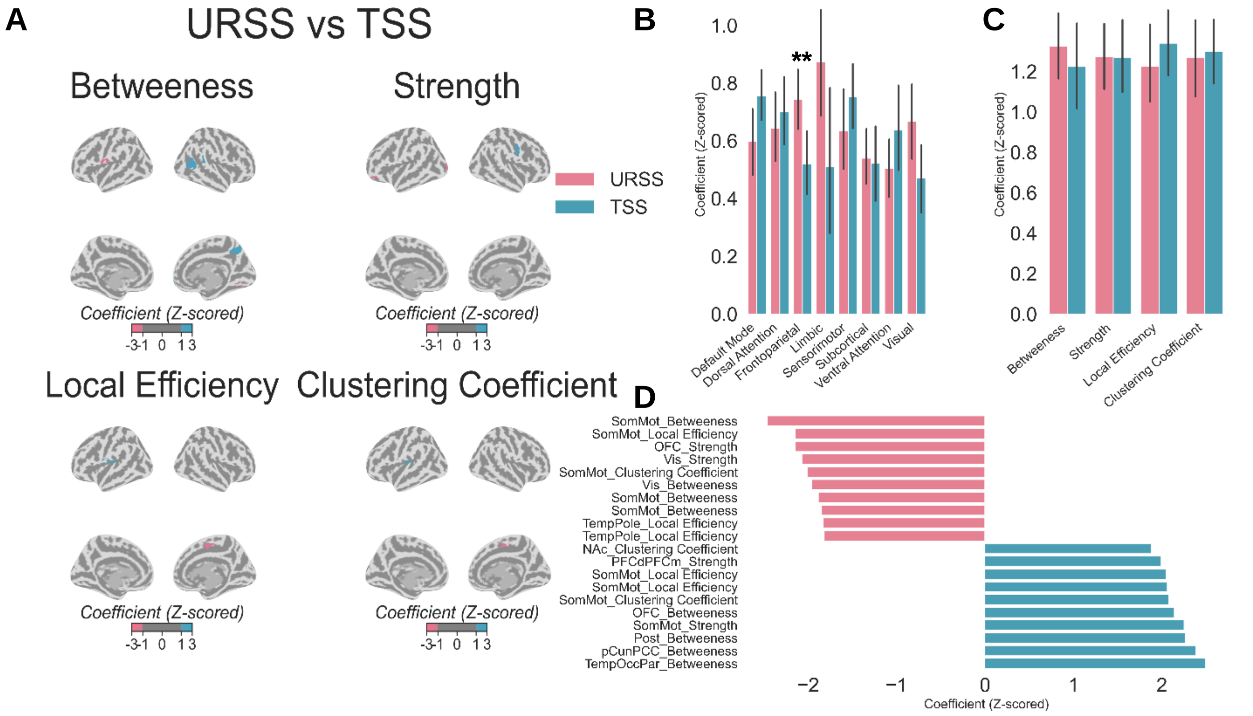

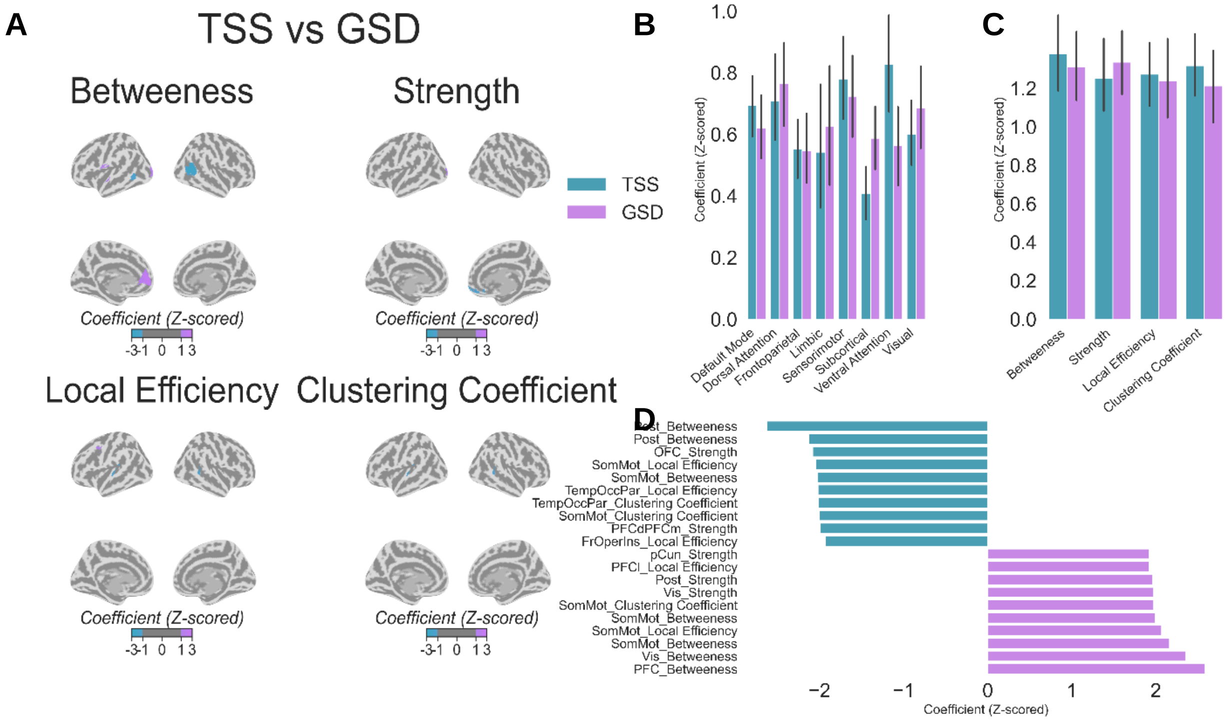


**Supplementary Figure 2.** (A) Surface maps illustrating the statistically significant coefficient differences between TSS and the remaining phenotypes. Regions highlighted on the brain differ significantly between the two groups, with the direction of the difference indicated by the colouring. (B) Bar plots displaying the average coefficient difference for each network across the two phenotypes, with vertical lines representing 95% confidence intervals. (C) Bar plots showing the average coefficient for each graph theory measure for both phenotypes, with vertical lines representing 95% confidence intervals. (D) Bar plots highlighting the top ten region and graph theory measure combinations with the largest coefficients for each phenotype. Abbreviations: taste and smell sensitivities (TSS); generalized sensory difficulties (GSD); movement difficulties and low energy (MDLE); under-responsive and sensory seeking (URSS). Temporal (Temp); Orbitofrontal Cortex (OFC); Prefrontal Cortex (PFC); Posterior (Post); Parietal (Par); Posterior Putamen (pPUT); Frontal Eye Fields (FEF); Somatomotor (SomMot); Visual Cortex (Vis); Frontal Operculum Insula (FrOperIns); Parahippocampal Cortex (PHC); Thalamus (THA); Temporal-Occipital-Parietal (TempOccPar)

### Supplementary Tables

### Table 1S

*Highly Influential Features for the SA vs GSD comparison*

**SA vs GSD**

| Phenotype | Region | Measure | Network | Coefficient |
|---|---|---|---|---|
| GSD | 7Networks_LH_SomMot_8 | Betweeness | SMS | 2.057895 |
| SA | 7Networks_RH_Default_Temp_1 | Strength | DMN | 2.448375 |
| SA | 7Networks_RH_Default_PFCdPFCm_1 | Strength | DMN | 2.346061 |
| SA | 7Networks_RH_Limbic_OFC_3 | Strength | Limbic | 2.328873 |
| SA | 7Networks_LH_Cont_PFCl_1 | Strength | FPN | 2.003824 |
| SA | 7Networks_LH_Default_PFC_4 | Strength | DMN | 1.955468 |
| SA | 7Networks_LH_Default_PFC_3 | Strength | DMN | 1.951869 |
| GSD | 7Networks_LH_DorsAttn_Post_8 | Local Efficiency | DA | 1.968821 |
| GSD | 7Networks_RH_SomMot_14 | Local Efficiency | SMS | 2.03015 |
| GSD | 7Networks_RH_DorsAttn_FEF_2 | Local Efficiency | DA | 2.417356 |
| SA | 7Networks_LH_Vis_4 | Clustering Coefficient | Visual | 1.973713 |
| GSD | 7Networks_LH_DorsAttn_Post_8 | Clustering Coefficient | DA | 1.944111 |
| GSD | 7Networks_RH_SomMot_14 | Clustering Coefficient | SMS | 2.039652 |
| GSD | 7Networks_RH_DorsAttn_FEF_2 | Clustering Coefficient | DA | 2.296409 |

### Table 2S

*Highly Influential Features for the SA vs MDLE comparison*

**SA vs MDLE**

| Phenotype | Region | Measure | Network | Coefficient |
|---|---|---|---|---|
| SA | 7Networks_RH_DorsAttn_Post_4 | Betweeness | DA | 1.992046 |
| MDLE | 7Networks_LH_Default_PHC_1 | Betweeness | DMN | 2.000876 |
| MDLE | 7Networks_RH_Limbic_TempPole_3 | Betweeness | Limbic | 2.050189 |
| MDLE | 7Networks_RH_Default_Par_2 | Betweeness | DMN | 2.260867 |
| SA | 7Networks_RH_Limbic_OFC_3 | Strength | Limbic | 2.453504 |
| SA | 7Networks_LH_Default_PFC_4 | Strength | DMN | 2.241101 |
| SA | 7Networks_LH_SomMot_9 | Strength | SMS | 2.146714 |
| SA | 7Networks_RH_Default_Temp_1 | Strength | DMN | 2.013338 |
| MDLE | 7Networks_RH_DorsAttn_Post_10 | Strength | DA | 1.989892 |
| MDLE | TianSubcortexS2_THA_DP_rh | Strength | SubCort | 2.091382 |
| MDLE | 7Networks_LH_SomMot_14 | Strength | SMS | 2.386036 |

| | | | | |
|---|---|---|---|---|
| SA | 7Networks_RH_SalVentAttn_FrOperIns_1 | Local Efficiency | VA | 2.172442 |
| MDLE | 7Networks_RH_DorsAttn_Post_4 | Local Efficiency | DA | 2.363352 |
| MDLE | 7Networks_RH_DorsAttn_Post_4 | Clustering Coefficient | DA | 2.162145 |

### Table 3S

*Highly Influential Features for the GSD vs URSS comparison*

**URSS vs GSD**

| Phenotype | Region | Measure | Network | Coefficient |
|---|---|---|---|---|
| *URSS* | 7Networks_LH_Vis_14 | Betweeness | Visual | 2.623285 |
| *URSS* | 7Networks_LH_SalVentAttn_Med_3 | Betweeness | VA | 2.293753 |
| *URSS* | 7Networks_LH_DorsAttn_PrCv_1 | Betweeness | DA | 2.105787 |
| *URSS* | 7Networks_LH_Default_pCunPCC_4 | Betweeness | DMN | 2.047801 |
| *GSD* | 7Networks_RH_Default_Temp_4 | Betweeness | DMN | 2.218307 |
| *URSS* | 7Networks_LH_Cont_OFC_1 | Strength | FPN | 2.139336 |
| *GSD* | 7Networks_RH_Default_PFCdPFCm_1 | Local Efficiency | DMN | 1.98809 |
| *GSD* | 7Networks_RH_SomMot_16 | Clustering Coefficient | SMS | 1.984739 |
| *GSD* | TianSubcortexS2_THA_VP_rh | Clustering Coefficient | SubCort | 2.001987 |
| *GSD* | TianSubcortexS2_NAc_core_rh | Clustering Coefficient | SubCort | 2.1252 |

### Table 4S

*Highly Influential Features for the TSS vs URSS comparison*

**TSS vs URSS**

| Phenotype | Region | Measure | Network | Coefficient |
|---|---|---|---|---|
| URSS | 7Networks_LH_SomMot_4 | Betweeness | SMS | 2.460834 |
| URSS | 7Networks_RH_Vis_4 | Betweeness | Visual | 1.958105 |
| TSS | 7Networks_RH_Limbic_OFC_3 | Betweeness | Limbic | 2.141546 |
| TSS | 7Networks_RH_DorsAttn_Post_2 | Betweeness | DA | 2.265359 |
| TSS | 7Networks_RH_Default_pCunPCC_3 | Betweeness | DMN | 2.381955 |
| TSS | 7Networks_RH_SalVentAttn_TempOccPar_2 | Betweeness | VA | 2.492399 |
| URSS | 7Networks_LH_Cont_OFC_1 | Strength | FPN | 2.143057 |
| URSS | 7Networks_LH_Vis_9 | Strength | Visual | 2.073048 |
| TSS | 7Networks_RH_Default_PFCdPFCm_5 | Strength | DMN | 1.993645 |
| TSS | 7Networks_RH_SomMot_7 | Strength | SMS | 2.250891 |

| | | | | |
|---|---|---|---|---|
| **URSS** | **7Networks_RH_SomMot_11** | **Local Efficiency** | **SMS** | **2.1463** |
| **TSS** | **7Networks_LH_SomMot_3** | **Local Efficiency** | **SMS** | **2.046436** |
| **TSS** | **7Networks_LH_SomMot_4** | **Local Efficiency** | **SMS** | **2.062891** |
| **URSS** | **7Networks_RH_SomMot_11** | **Clustering Coefficient** | **SMS** | **2.013165** |
| **TSS** | **7Networks_LH_SomMot_3** | **Clustering Coefficient** | **SMS** | **2.078573** |

### Table 5S

*Highly Influential Features for the TSS vs SA comparison*

**TSS vs SA**

| Phenotype | Region | Measure | Network | Coefficient |
|---|---|---|---|---|
| ***SA*** | **7Networks_LH_Default_PFC_6** | **Betweeness** | **DMN** | **2.792324** |
| ***SA*** | **7Networks_LH_SomMot_4** | **Betweeness** | **SMS** | **2.275429** |
| ***SA*** | **7Networks_LH_Cont_OFC_1** | **Betweeness** | **FPN** | **2.06478** |
| ***TSS*** | **7Networks_RH_Default_Temp_3** | **Betweeness** | **DMN** | **1.955676** |
| ***TSS*** | **7Networks_RH_Default_pCunPCC_3** | **Betweeness** | **DMN** | **2.029544** |
| ***TSS*** | **7Networks_RH_Default_PFCdPFCm_7** | **Betweeness** | **DMN** | **2.171744** |
| ***TSS*** | **7Networks_RH_DorsAttn_Post_2** | **Betweeness** | **DA** | **2.220726** |
| ***SA*** | **7Networks_LH_Cont_PFCl_1** | **Strength** | **FPN** | **2.147591** |
| ***SA*** | **7Networks_RH_Default_pCunPCC_1** | **Strength** | **DMN** | **2.065794** |
| ***SA*** | **7Networks_RH_SomMot_11** | **Local Efficiency** | **SMS** | **2.264713** |
| ***TSS*** | **7Networks_LH_Default_PFC_6** | **Local Efficiency** | **DMN** | **1.968399** |
| ***TSS*** | **7Networks_RH_DorsAttn_Post_4** | **Local Efficiency** | **DA** | **1.971887** |
| ***SA*** | **7Networks_RH_SomMot_11** | **Clustering Coefficient** | **SMS** | **2.294636** |
| ***TSS*** | **7Networks_LH_Default_PFC_6** | **Clustering Coefficient** | **DMN** | **1.936566** |
| ***TSS*** | **7Networks_RH_DorsAttn_Post_4** | **Clustering Coefficient** | **DA** | **1.971482** |

### Table 6S

*Highly Influential Features for the TSS vs MDLE comparison*

**MDLE vs TSS**

| Phenotype | Region | Measure | Network | Coefficient |
|---|---|---|---|---|
| *MDLE* | TianSubcortexS2_pPUT_lh | Betweeness | SubCort | 2.354358 |
| *MDLE* | 7Networks_LH_Default_PFC_6 | Betweeness | DMN | 2.295117 |
| *MDLE* | 7Networks_LH_SomMot_4 | Betweeness | SMS | 2.006512 |
| *MDLE* | 7Networks_RH_Limbic_TempPole_3 | Betweeness | Limbic | 1.987032 |
| *TSS* | 7Networks_RH_SalVentAttn_Med_2 | Betweeness | VA | 2.012489 |
| *TSS* | 7Networks_RH_Default_PFCdPFCm_7 | Betweeness | DMN | 2.0786 |
| *TSS* | 7Networks_RH_Limbic_OFC_3 | Betweeness | Limbic | 2.120717 |
| *TSS* | 7Networks_RH_DorsAttn_Post_2 | Betweeness | DA | 2.172222 |
| *TSS* | 7Networks_RH_Default_pCunPCC_3 | Betweeness | DMN | 2.257454 |
| *TSS* | 7Networks_RH_SalVentAttn_TempOccPar_2 | Betweeness | VA | 2.51792 |
| *MDLE* | 7Networks_LH_Vis_9 | Strength | Visual | 2.048107 |
| *TSS* | 7Networks_RH_Limbic_OFC_3 | Strength | Limbic | 2.298453 |
| *TSS* | 7Networks_RH_SomMot_7 | Strength | SMS | 2.527625 |
| *TSS* | 7Networks_LH_SomMot_4 | Local Efficiency | SMS | 1.959909 |
| *TSS* | 7Networks_RH_SomMot_5 | Local Efficiency | SMS | 1.995625 |
| *TSS* | 7Networks_LH_Default_PFC_6 | Local Efficiency | DMN | 2.180638 |
| *TSS* | 7Networks_RH_SomMot_5 | Clustering Coefficient | SMS | 2.016443 |
| *TSS* | 7Networks_LH_Default_PFC_6 | Clustering Coefficient | DMN | 2.136822 |

### Table 7S

*Highly Influential Features for the TSS vs GSD comparison*

**GSD vs TSS**

| Phenotype | Region | Measure | Network | Coefficient |
|---|---|---|---|---|
| *TSS* | 7Networks_RH_DorsAttn_Post_2 | Betweeness | DA | 2.622994 |
| *TSS* | 7Networks_LH_DorsAttn_Post_2 | Betweeness | DA | 2.127163 |
| *TSS* | 7Networks_RH_SomMot_14 | Betweeness | SMS | 2.021785 |
| *GSD* | 7Networks_LH_SomMot_1 | Betweeness | SMS | 1.992794 |
| *GSD* | 7Networks_LH_SomMot_4 | Betweeness | SMS | 2.163798 |
| *GSD* | 7Networks_LH_Vis_9 | Betweeness | Visual | 2.3593 |
| *GSD* | 7Networks_LH_Default_PFC_6 | Betweeness | DMN | 2.584466 |
| *TSS* | 7Networks_RH_Limbic_OFC_3 | Strength | Limbic | 2.081115 |
| *TSS* | 7Networks_RH_Default_PFCdPFCm_1 | Strength | DMN | 1.99333 |
| *GSD* | 7Networks_LH_DorsAttn_Post_5 | Strength | DA | 1.963456 |
| *GSD* | 7Networks_LH_Vis_9 | Strength | Visual | 1.972035 |
| *TSS* | 7Networks_LH_SomMot_2 | Local Efficiency | SMS | 2.041564 |
| *TSS* | 7Networks_RH_SalVentAttn_TempOccPar_1 | Local Efficiency | VA | 2.018622 |
| *GSD* | 7Networks_LH_Cont_PFCl_5 | Local Efficiency | FPN | 1.923877 |
| *GSD* | 7Networks_RH_SomMot_14 | Local Efficiency | SMS | 2.069318 |
| *TSS* | 7Networks_RH_SalVentAttn_TempOccPar_1 | Clustering Coefficient | VA | 2.01487 |

| | | | | |
|---|---|---|---|---|
| *TSS* | 7Networks_LH_SomMot_2 | Clustering Coefficient | SMS | 2.00468 |
| *GSD* | 7Networks_RH_SomMot_14 | Clustering Coefficient | SMS | 1.975759 |
